# Individual differences in reading comprehension and off-task thought relate to perceptually-coupled and decoupled cognition

**DOI:** 10.1101/387035

**Authors:** Meichao Zhang, Nicola Savill, Daniel S. Margulies, Jonathan Smallwood, Elizabeth Jefferies

**Author notes:** To whom correspondence should be addressed. Department of Psychology, the University of York, Heslington, York, UK. Email address (M. Zhang). Email addresses of all authors (M. Zhang), (N. Savill), (D. Margulies), (J. Smallwood), (E. Jefferies).

## Abstract

Although the default mode network (DMN) is associated with off-task states, recent evidence shows it can support tasks. This raises the question of how DMN activity can be both beneficial and detrimental to task performance. The decoupling hypothesis proposes that these opposing states occur because DMN supports modes of cognition driven by external input, as well as retrieval states unrelated to input. To test this account, we capitalised on the fact that during reading, regions in DMN are thought to represent the meaning of words through their coupling with visual cortex; the absence of visual coupling should occur when the attention drifts off from the text. We examined individual differences in reading comprehension and off-task thought while participants read an expository text in the laboratory, and related variation in these measures to (i) the neural response during reading in the scanner (Experiment 1), and (ii) patterns of intrinsic connectivity measured in the absence of a task (Experiment 2). The responsiveness of a region of DMN in middle temporal gyrus (MTG) to orthographic inputs during reading predicted good comprehension, while intrinsic decoupling of the same site from visual cortex at rest predicted more frequent off-task thought. In addition, good comprehension was associated with greater intrinsic connectivity between MTG and medial prefrontal regions also within DMN, demonstrating that DMN coupling can support task performance, not only off-task states. These findings indicate that the opposing roles of DMN in cognition reflect its capacity to support both perceptually-coupled and decoupled cognition.

## 1. Introduction

The default mode network (DMN) is implicated in off-task lapses (Christoff, Gordon, Smallwood, Smith, & Schooler, 2009; Mason et al., 2007; Stawarczyk, Majerus, Maquet, & D’Argembeau, 2011), when its activation can be linked to poor performance in an ongoing task. However, more recently, DMN has been shown to support demanding tasks such as comprehension (Smallwood et al., 2013), semantic decisions (Krieger-Redwood et al., 2016; Murphy et al., 2018), and working memory (Vatansever, Menon, Manktelow, Sahakian, & Stamatakis, 2015). A key puzzle therefore concerns how regions within the DMN can support these conflicting states of good and poor performance. Theories of DMN function emphasise that this network is situated far along an information-processing hierarchy (Margulies et al., 2016; Vidaurre, Smith, & Woolrich, 2017), allowing the integration of diverse features and the abstraction of knowledge from experience in the physical world (Lambon Ralph, Jefferies, Patterson, & Rogers, 2017; Patterson, Nestor, & Rogers, 2007). The distance of DMN from sensory-motor regions might allow DMN regions to support ‘perceptually decoupled’ states such as off-task thought. However, highly-integrated representations might also be necessary for externally-presented tasks supported by the DMN. This gives rise to the hypothesis that different patterns of connectivity from the same DMN regions might be responsible for conflicting states such as (i) good comprehension, when DMN regions in the semantic network are focused on extracting meaning from external input and (ii) poor comprehension, following off-task lapses, when the same DMN regions might be perceptually-decoupled. A better understanding of how DMN can contribute to these opposing states might therefore come from studies of individual differences in comprehension, whilst off-task lapses within the comprehension task are recorded.

The ventral visual stream, which supports visual-to-semantic processing within the temporal lobe, is implicated in reading comprehension (Dehaene & Cohen, 2011; Dehaene, Cohen, Morais, & Kolinsky, 2015; Spitsyna, Warren, Scott, Turkheimer, & Wise, 2006). Neural regions along the posterior-to-anterior axis of the temporal lobe increasingly respond to meaning as opposed to visual properties (Gauthier, Eger, Hesselmann, Giraud, & Kleinschmidt, 2012; Patterson et al., 2007; Spitsyna et al., 2006). The lateral anterior temporal lobes (ATL), at the top of this hierarchy, are thought to represent heteromodal concepts, following the convergence of meaningful inputs across modalities (Lambon Ralph et al., 2017; Murphy et al., 2017; Visser, Jefferies, & Lambon Ralph, 2010). Nevertheless, the same semantic representations might play a role in spontaneous thought.

Variation in connectivity from ATL predicts individual differences in patterns of spontaneous thought (Smallwood et al., 2016; Vatansever et al., 2017). During self-generated thought, semantic representations might become “perceptually decoupled” – i.e., they may no longer be driven by inputs from the external world. Individual differences in the extent to which semantic cognition is perceptually-coupled (for example, focussed on the text to be understood), or perceptually decoupled (resulting in off-task lapses) might give rise to the previously-reported negative correlation between comprehension and off-task thought during reading (Sanders, Wang, Schooler, & Smallwood, 2017; Smallwood, McSpadden, & Schooler, 2008).

To test this hypothesis, we examined individual differences in comprehension and off-task thought during reading in two experiments where the task context offers us the opportunity to gain an understanding of these opposing roles of the DMN. Experiment 1 employed an online sentence reading task during fMRI: we examined differences in activation that could predict comprehension outside the scanner, to test the hypothesis that people who show greater perceptual coupling within semantic regions in the DMN will have better comprehension of externally-presented material. In Experiment 2, we examined individual differences in intrinsic connectivity at rest from this same DMN region to establish how different patterns of coupling might give rise to excellent understanding of text in some people and inattentiveness and off-task thought in others.

## 2. Materials and Methods

We measured reading comprehension outside the scanner and assessed (i) comprehension of the text; (ii) how frequently people noticed that they had stopped paying attention to the meaning of the text during reading and (iii) what people reported they had been thinking about after they had completed the reading task. We performed two fMRI experiments with this cohort. In Experiment 1 (task-based fMRI reading task), participants passively-viewed sentences and nonwords in the scanner, to assess their responsiveness to meaningful and non-meaningful visual information. To anticipate, we found a region in lateral temporal cortex extending into DMN that showed a stronger response to visually-presented material in good comprehenders. In Experiment 2 (resting-state fMRI), we then related individual differences in the intrinsic connectivity of this region to comprehension and off-task thought, to test the hypothesis that different patterns of connectivity to visual and DMN predict performance.

### 2.1 Participants

Ethical approval was obtained from the Department of Psychology and York Neuroimaging Centre, University of York ethics committees. Sixty-nine undergraduate or postgraduate students were recruited for this study (age range 18-31, mean age = 19.87 ± 2.33, 26 males). All were right-handed native English speakers, and had normal or corrected-to-normal vision. None of them had any history of neurological impairment, diagnosis of learning difficulty or psychiatric illness. All provided written informed consent prior to taking part and received a monetary reward for their participation.

### 2.2 Behavioural assessment

*Off-task frequency and comprehension*. Following an MRI scan (details below), participants were asked to complete a battery of behavioural assessments examining reading comprehension and off-task thought. In order to create a naturalistic reading experience, we presented the text on a printed booklet. The text was selected and shortened from Bill Bryson’s “A short story of everything” (Smallwood et al., 2013), in font size 14, 1.5 line spacing. The passage was about the topic of geology (word count = 1050). During reading, the participants were required to note down any moments when they noticed they had stopped paying attention to the meaning of the text, by circling the word they had reached at this point. A detailed instruction booklet was used to guide the participants through the experiment. After they finished reading, they were asked to answer 17 open-ended questions assessing their comprehension of what they read, without being able to refer back to the text (Smallwood et al., 2013). The answers to the comprehension questions were scored for accuracy referring to the standard answers by two experimenters. Responses were given a score of 1 if they contained key information, and otherwise a score of 0. The two scorers produced very similar ratings (*r* = .92, *p* < .001). The experiment took approximately 30 (±5) minutes.

*Off-task experiences*. A self-report measurement, the New-York Cognition Questionnaire (NYC-Q), was also used to assess off-task behaviour during the reading task. The first section contained 22 questions about the content of thoughts (e.g., *I thought about personal worries*), rated on a scale of 1 (*Completely did not describe my thoughts*) to 9 (*Completely did describe my thoughts*). The second section contained 8 questions about the form of these thoughts (e.g., *whilst I was reading my thoughts were in the form of images*), rated on a scale of 1 (*Completely did not characterize my experience*) to 9 (*Completely did characterize my experience*; Gorgolewski et al., 2014; Sanders et al., 2017). In the current study, we limited our analysis to the 22 questions relating to the content of off-task thought. We calculated an overall average for each participant, which is thought to reflect how much each individual was being off-task. In this way, we assessed both off-task frequency (i.e., moments in which attention was not directed towards the reading task) and the nature of these experiences (the extent to which coherent internally-generated thoughts predominated). Theories have suggested these measures are not identical: off-task moments are necessary but not sufficient for mind-wandering, which are characterised by a pattern of sustained retrieval from memory (e.g., of autobiographical episodes; Smallwood, 2013). Prior to data analysis, outliers more than 2.5 standard deviations above or below the mean were imputed with these cut-off values and all variables were z-transformed.

### 2.3 Neuroimaging data acquisition

Structural and functional data were acquired using a 3T GE HDx Excite Magnetic Resonance Imaging (MRI) scanner utilizing an eight-channel phased array head coil at the York Neuroimaging Centre, University of York. Structural MRI acquisition in all participants was based on a T1-weighted 3D fast spoiled gradient echo sequence (repetition time (TR) = 7.8 s, echo time (TE) = minimum full, flip angle = 20°, matrix size = 256 × 256, 176 slices, voxel size = 1.13 mm × 1.13 mm × 1 mm).

The reading task in Experiment 1 used single-shot 2D gradient-echo-planar imaging (TR = 3 s, TE = minimum full, flip angle = 90°, matrix size = 64 × 64, 60 slices, voxel size = 3 mm × 3 mm × 3 mm, 80 volumes). The participants passively viewed meaningful sentences (e.g., her + secrets + were + written + in + her + diary) and meaningless sequences of non-words (e.g., crark + dof + toin + mesk + int + lisal + glod + flid), item-by-item. In total, there were 10 meaningful sentences, taken from Rodd, Davis, and Johnsrude (2005), and 10 nonword lists, matched for both word length and number of syllables. Word and nonword sets were each presented in two blocks in a pseudo-random order (i.e., a total of 4 blocks). A task instruction (e.g., Meaningful) was used to indicate the transition between different conditions. Each sequence ended with a red fixation lasting 4000-6000 ms. Each word or nonword was presented for 600 ms, followed by a 250 ms fixation before the next item was presented. A fluid-attenuated inversion-recovery (FLAIR) scan with the same orientation as the functional scans was collected to improve co-registration between subject-specific structural and functional scans.

A 9-minute resting-state fMRI scan was used in Experiment 2, recorded using single-shot 2D gradient-echo-planar imaging (TR = 3 s, TE = minimum full, flip angle = 90°, matrix size = 64 × 64, 60 slices, voxel size = 3 mm × 3 mm × 3 mm, 180 volumes). During resting-state scanning, the participants were instructed to focus on a fixation cross with their eyes open and to keep as still as possible, without thinking about anything in particular. Neuroimaging data for Experiments 1 and 2 were collected in the same session, with the resting-state sequence presented first.

### 2.4 Neuroimaging data pre-processing

All functional and structural data were pre-processed using a standard pipeline and analysed via the FMRIB Software Library (FSL version 6.0, www.fmrib.ox.ac.uk/fsl). Individual FLAIR and T1-weighted structural brain images were extracted using FSL’s Brain Extraction Tool (BET). Structural images were linearly registered to the MNI152 template using FMRIB’s Linear Image Registration Tool (FLIRT). The reading functional neuroimaging data were pre-processed and analysed by using FSL’s FMRI Expert Analysis Tool (FEAT). A standard pre-processing pipeline was applied, including motion correction via MCFLIRT, slice-timing correction using Fourier space time-series phase-shifting, and spatial smoothing using a Gaussian kernel of FWHM 6 mm. In addition, for the task-based fMRI data in Experiment 1, high-pass temporal filtering (sigma = 100 s) was applied in order to remove temporal signal drift. For the resting-state fMRI data in Experiment 2, both high pass (sigma = 200 s) and low-pass temporal filtering (sigma = 2.8 s) was applied, in order to constrain analyses to low-frequency fluctuations.

### 2.5 Neuroimaging analysis

#### 2.5.1 Task-based fMRI analysis (Experiment 1)

This analysis identified sites in which activation during the reading task was modulated by comprehension, off-task frequency or the content of off-task thought (i.e., NYC-Q). In the first-level analysis of the reading task performed in the scanner, we identified voxels responding to (i) meaning and (ii) orthographic inputs devoid of meaning, through the contrasts of *Meaningful > Baseline*, *Meaningless > Baseline*, *Meaningful > Meaningless*, plus the reverse, for each participant. In the higher-level analysis at the group level, z-transformed behavioural data for comprehension, off-task frequency and NYC-Q were added as explanatory variables, using FMRIB’s Local Analysis of Mixed Effects (FLAME1), with automatic outlier de-weighting (Woolrich, 2008). A 50% probabilistic grey-matter mask was applied. Clusters were thresholded using Gaussian random-field theory, with a cluster-forming threshold of *z* = 2.6 and a familywise-error-corrected significance level of *p* = .05.

#### 2.5.2 Resting-state fMRI analysis (Experiment 2)

We next considered whether the intrinsic connectivity of regions identified in Experiment 1 predicted reading comprehension and off-task thought. A cluster which showed a stronger response to orthographic input in people with good comprehension overlapped with both the default mode network (DMN), which is typically task-negative, and the adjacent fronto-partietal network (FPN), which is typically task-positive. In the literature, both networks are implicated in semantic cognition (Davey et al., 2016). We therefore masked the results of Experiment 1 by these DMN and FPN networks, defined by a parcellation of 1000 resting-state scans (Yeo et al., 2011), obtained from Freesurfer (https://surfer.nmr.mgh.harvard.edu/fswiki/CorticalParcellation_Yeo2011). This identified a region of middle temporal gyrus (MTG) within DMN, and a region of inferior temporal gyrus (ITG) in the FPN. These regions were taken as seeds in a subsequent analysis of intrinsic connectivity (see Figure 2B).

**Fig. 2.**
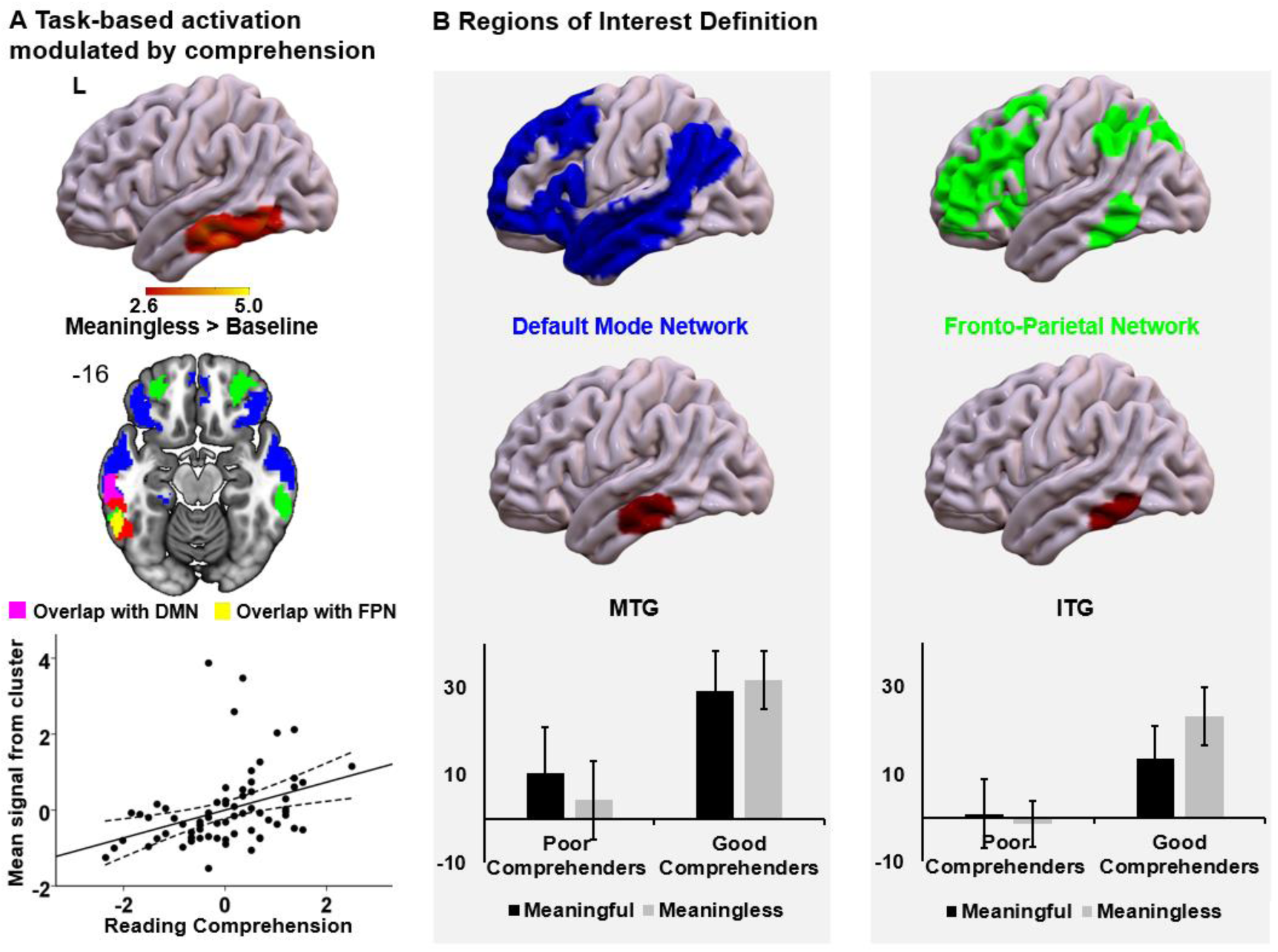
A) Regions modulated by comprehension in task-based fMRI (Experiment 1). Participants were asked to passively read meaningful sentences and meaningless sequences of non-words. The image shows voxels that were more active for good comprehenders for the contrast of meaningless nonwords over baseline (*z* > 2.6; *p* < .05; in red). This cluster overlaps with two large-scale brain networks previously implicated in semantic cognition: default mode network (DMN; in blue) and fronto-parietal network (FPN; in green). These networks were defined by Yeo et al. (2011), in a 7-network parcellation of whole-brain functional connectivity for 1000 brains. The network map is fully saturated to emphasize the regions of overlap (in pink and yellow). The number in the top left of the overlap map indicates the coordinate value of the corresponding plane. The scatterplot presents the correlation between the mean signal extracted from the significant cluster and reading comprehension. The error lines on the scatterplot indicate the 95% confidence estimates of the mean. Each point describes an individual participant. L = Left. **B) Definition of task-based ROIs.** The seeds used for resting-state functional connectivity in Experiment 2 were defined as the overlaps between the region in which task-based activation to meaningless orthographic input was modulated by comprehension and two large-scale brain networks: DMN and FPN. This gave rise to two ROIs: middle temporal gyrus (MTG) in DMN and inferior temporal gyrus (ITG) in FPN. The bar charts in the lower panel present the average of mean signal extracted from MTG and ITG in the contrasts of *Meaningful > Baseline* and *Meaningless > Baseline* for poor and good comprehenders. The error bars indicate the standard error in each contrast in each group.

We extracted the time series from the seeds and used this data as explanatory variables in whole-brain connectivity analyses at the single subject level. Sixty-four participants were included in this analysis (five participants without intrinsic connectivity data were excluded). These functional connectivity maps were then related to individual differences in behaviour using a multiple regression model, in which z-transformed scores for comprehension, off-task frequency and NYC-Q were added as explanatory variables. In order to control for the spurious correlations that might emerge from movement, we included two canonical components, group mean and mean framewise displacement (FD; Jenkinson, Bannister, Brady, & Smith, 2002), as nuisance covariates in the model. Automatic outlier de-weighting and a 50% probabilistic grey-matter mask were applied as before. Clusters were thresholded using Gaussian random-field theory, with a cluster-forming threshold of *z* = 2.6 and a familywise-error-corrected significance level of *p* = .05. We also applied Bonferroni correction to account for the fact that we included two models (IFG and MTG) and used two-tailed tests (in which behaviour could relate to both stronger and weaker connectivity). Consequently, the *p* value accepted as significant was *p* < .0125.

## 3. Results

### 3.1 Behavioural results

The behavioural results are summarized in Figure 1. Pearson’s correlation analysis revealed that off-task frequency was positively correlated with the NYC-Q scores (*r* = .51, *p* < .001). There was also a significant negative correlation between off-task frequency and comprehension scores (*r* = −.26, *p* = .029), suggesting that frequent off-task thought impairs reading comprehension, in line with previous findings (Sanders et al., 2017; Smallwood et al., 2008). However, the correlation between NYC-Q scores and comprehension scores was not significant (*r* = −.17, *p* = .16).

**Fig. 1.**
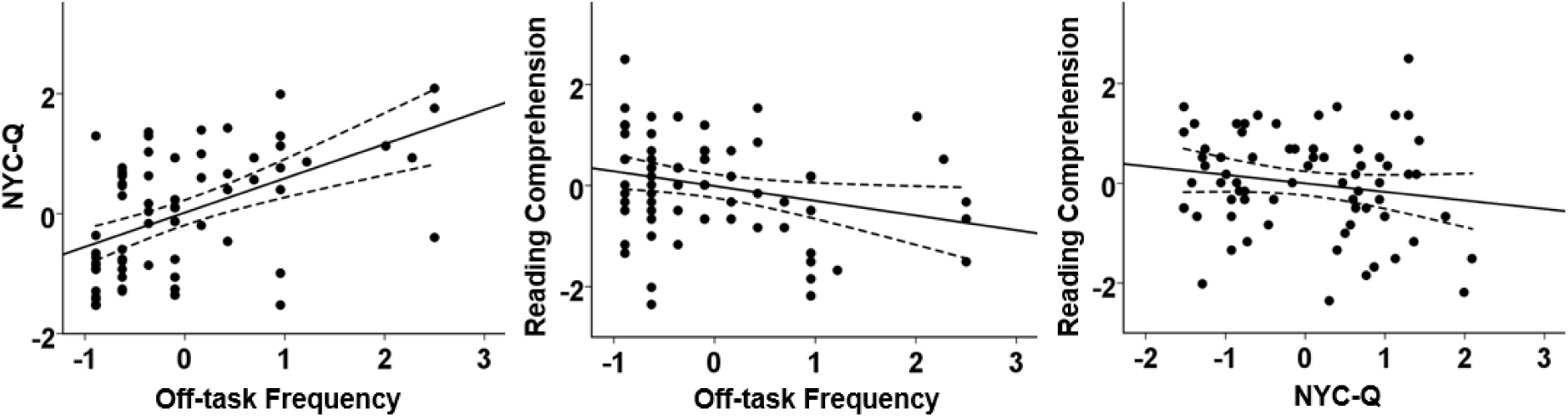
Behavioural results. The scatterplots present the correlations between off-task frequency, NYC-Q (i.e., the content of off-task thought), and reading comprehension. The error lines on the scatterplots indicate the 95% confidence estimates of the mean. Each point describes each participant.

### 3.2. Experiment 1: Reading task

A whole-brain analysis indicated that activation in middle/inferior temporal gyrus and temporal fusiform cortex in the contrast of *Meaningless > Baseline* was modulated by individual differences in reading comprehension (see Figure 2A). To understand the nature of this relationship, we plotted the relationship between mean signal in this region and reading comprehension across individuals. People with better comprehension showed a stronger response to orthographic input, even when this was not meaningful.

The cluster which showed a stronger response to orthographic input in people with good comprehension overlapped with both the default mode network (DMN; with overlap in pink), which is typically task-negative, and the adjacent fronto-partietal network (FPN; with overlap in yellow), which is typically task-positive (see Figure 2A). We identified voxels that showed greater activation to orthographic input and that also fell within DMN and FPN according to the network parcellation by Yeo et al. (2011; see Figure 2B). Within these DMN and FPN regions, we described individual differences in the responsiveness to meaningful and meaningless orthographic inputs relative to baseline, by performing a split-half analysis (i.e., the z-transformed score above or below 0) according to text comprehension scores outside the scanner. This showed that the effect of comprehension was not specific to nonwords; a similar pattern was seen for meaningful material. People with better comprehension showed a stronger response to orthographic inputs in both DMN and FPN-regions of the lateral temporal lobe, irrespective of meaning.

To more fully characterise the stronger BOLD response to orthographic input observed in participants with better comprehension, we compared this temporal lobe cluster (shown in red in Figure 3) with the main effects of meaning and orthographic input across the group. The effect of individual differences in comprehension lay adjacent to brain regions responsive to semantic processing (i.e., the contrast of *Meaningful > Meaningless*; shown in blue in Figure 3A). The cluster was also adjacent to and partly overlapping with brain regions that were responsive to orthographic inputs irrespective of meaning (i.e., the conjunction of the contrasts of *Meaningful > Baseline* and *Meaningless > Baseline*, shown in green in Figure 3B). This pattern is summarised in Figure 3C, which shows that the individual differences cluster (i.e. better comprehension associated with a stronger response to orthographic input) was located at the intersection of regions linked to (i) semantic processing and (ii) orthography. Such a region would be implicated in visual-to-semantic processes in contemporary accounts that propose graded abstraction from unimodal visual to heteromodal conceptual representations within the temporal lobe (Lambon Ralph et al., 2017).

**Fig. 3.**
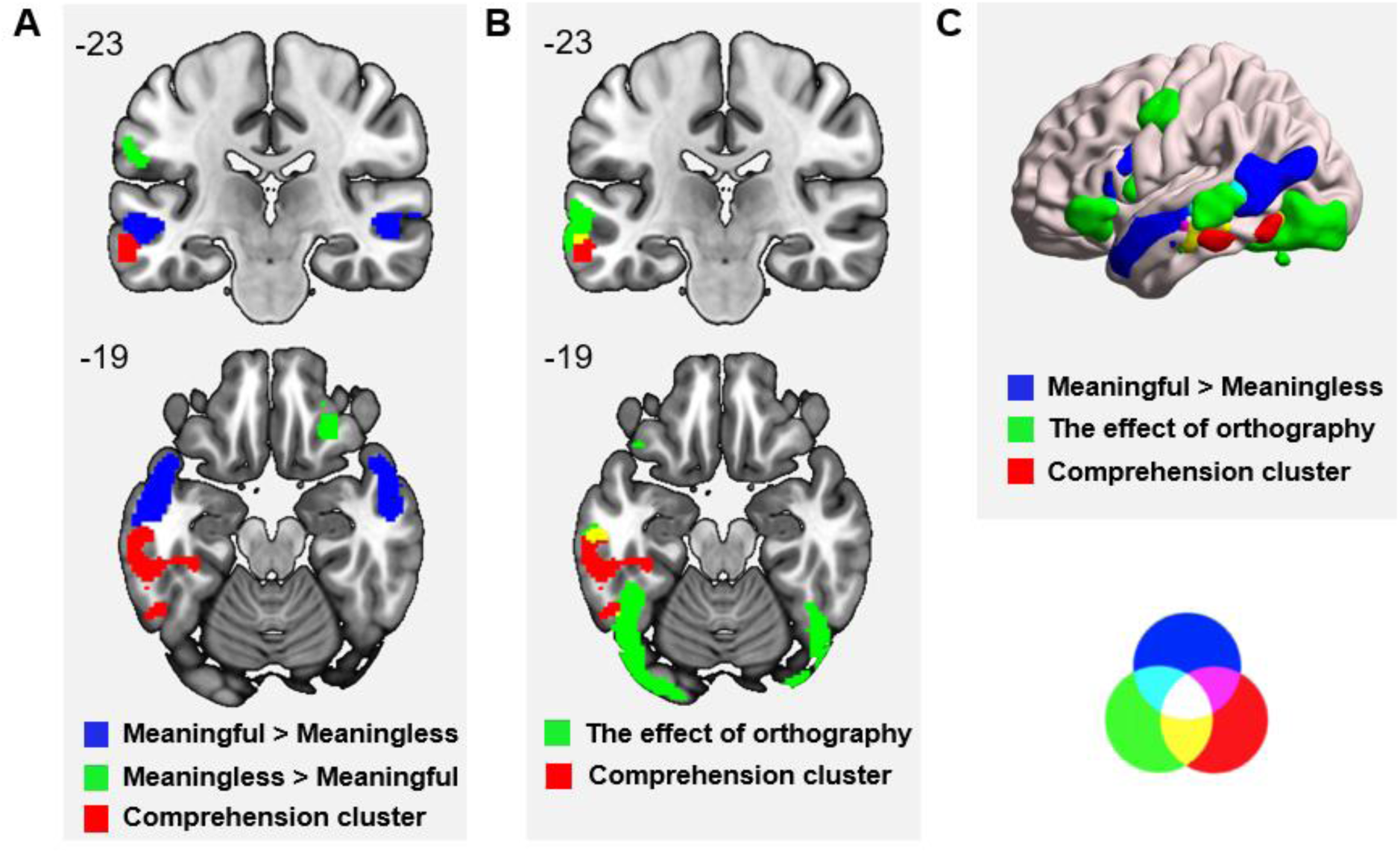
The overlap of the regions modulated by reading comprehension with **(A)** the activated regions in the contrasts of *Meaningful > Meaningless* and *Meaningless > Meaningful*, and **(B)** the response to orthographic input, as well as (**C)** the activated regions in the contrast of *Meaningful > Meaningless* and the activated regions respond to the effect of orthography. Regions in pink, yellow, blue, and white indicate regions of overlap. These maps are fully saturated to emphasize the regions of overlap. All maps are thresholded at *z* > 2.6 (*p* < .05). Numbers at the top left of each panel indicate the coordinate value of the corresponding plane. The overlapping circles at the lower right indicate the colour of overlap regions.

### 3.3. Experiment 2: Resting-state functional connectivity

Since our research focussed on the role of DMN in task-relevant and irrelevant patterns of semantic retrieval, we identified voxels that showed greater activation to orthographic input for good comprehenders in Experiment 1 and that also fell within the DMN according to a commonly-used whole-brain parcellation (Yeo et al., 2011). We used this DMN region, which fell within MTG, as a seed for an analysis of intrinsic connectivity in Experiment 2. For completeness, we also took voxels within the FPN as a second seed, producing an ROI focussed on ITG (see Figure 2B).

We explored whether individual differences in reading comprehension and off-task thought were associated with variation in patterns of intrinsic connectivity from these two seeds. We generated functional connectivity maps for each region, for each individual, and then analysed these spatial maps using a series of multiple regression analyses that included individual scores in off-task thought (i.e., off-task frequency and NYC-Q) and reading comprehension as explanatory variables. There were no significant differences in the connectivity of ITG that related to either off-task thought or comprehension (there were also no effects approaching significance that failed to survive Bonferroni correction for the number of seeds or the two-tailed nature of our tests). Consequently, this seed region is not discussed further below. However, as predicted, individual differences in the connectivity of MTG within the DMN did predict behaviour.

Group-level intrinsic connectivity maps for the MTG DMN seed region (i.e., irrespective of performance) are presented in Figure 4. To understand how the regions of positive and negative connectivity from this seed region correspond to the network implicated in semantic processing, we compared these spatial maps to a meta-analytic map generated for the term SEMANTIC using Neurosynth (Yarkoni, Poldrack, Nichols, Van Essen, & Wager, 2011). This revealed that regions of relatively high connectivity from MTG (shown in red in Figure 4) largely overlapped with regions important for semantic cognition (shown in green, with overlap in yellow).

**Fig. 4.**
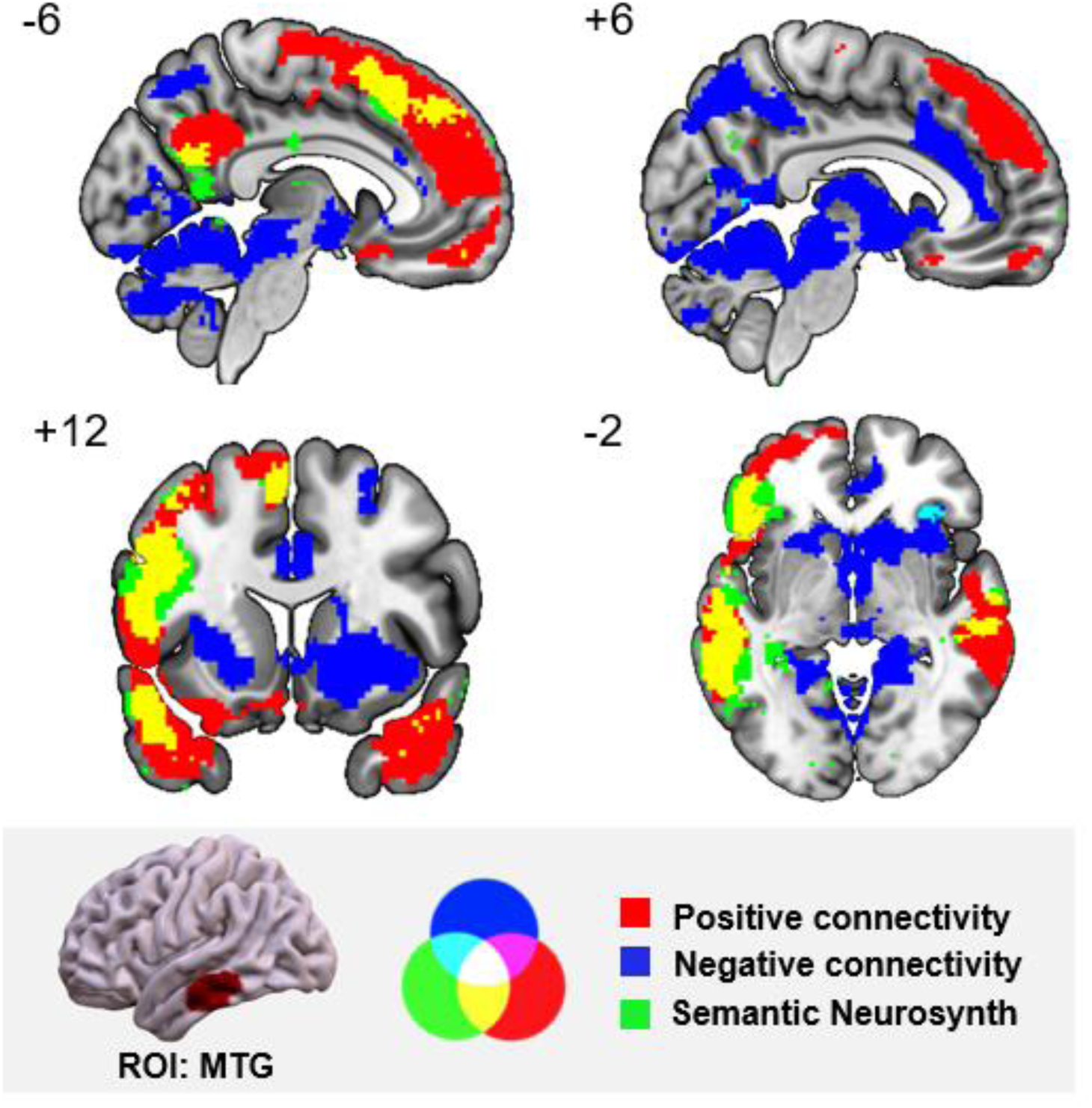
The group-level patterns of relatively high (in red) and low (in blue) functional connectivity from the DMN seed in MTG during resting-state fMRI (cluster correction, *z* > 2.6, *p* < .05), and the overlap of these positive and negative networks with a semantic meta-analytic map (regions in green) derived from Neurosynth (using ‘semantic’ as a search term). These maps are fully saturated to emphasize the regions of overlap. Numbers at the top left of each panel indicates the coordinate value of the corresponding plane. The bottom panel highlighted in grey shows the seed region and the overlapping circles that indicate the colour of overlap regions. MTG = middle temporal gyrus.

#### 3.3.1. Reading Comprehension

We found that MTG connectivity was modulated by reading comprehension. Participants with better comprehension scores showed stronger connectivity of the MTG DMN seed region to anterior cingulate cortex (cingulate gyrus and paracingulate gyrus). This cluster is illustrated in Figure 5. Of the voxels within the anterior cingulate cluster that fell within the large-scale networks defined by Yeo et al. (2011), 88.3% were within DMN, 11.3 % fell within FPN, and 0.4 % fell within ventral attention network. These findings show that connectivity between different nodes of DMN is not necessarily linked to more off-task thought and poorer performance; connectivity within DMN network also supports aspects of cognition relevant to comprehension.

**Fig. 5.**
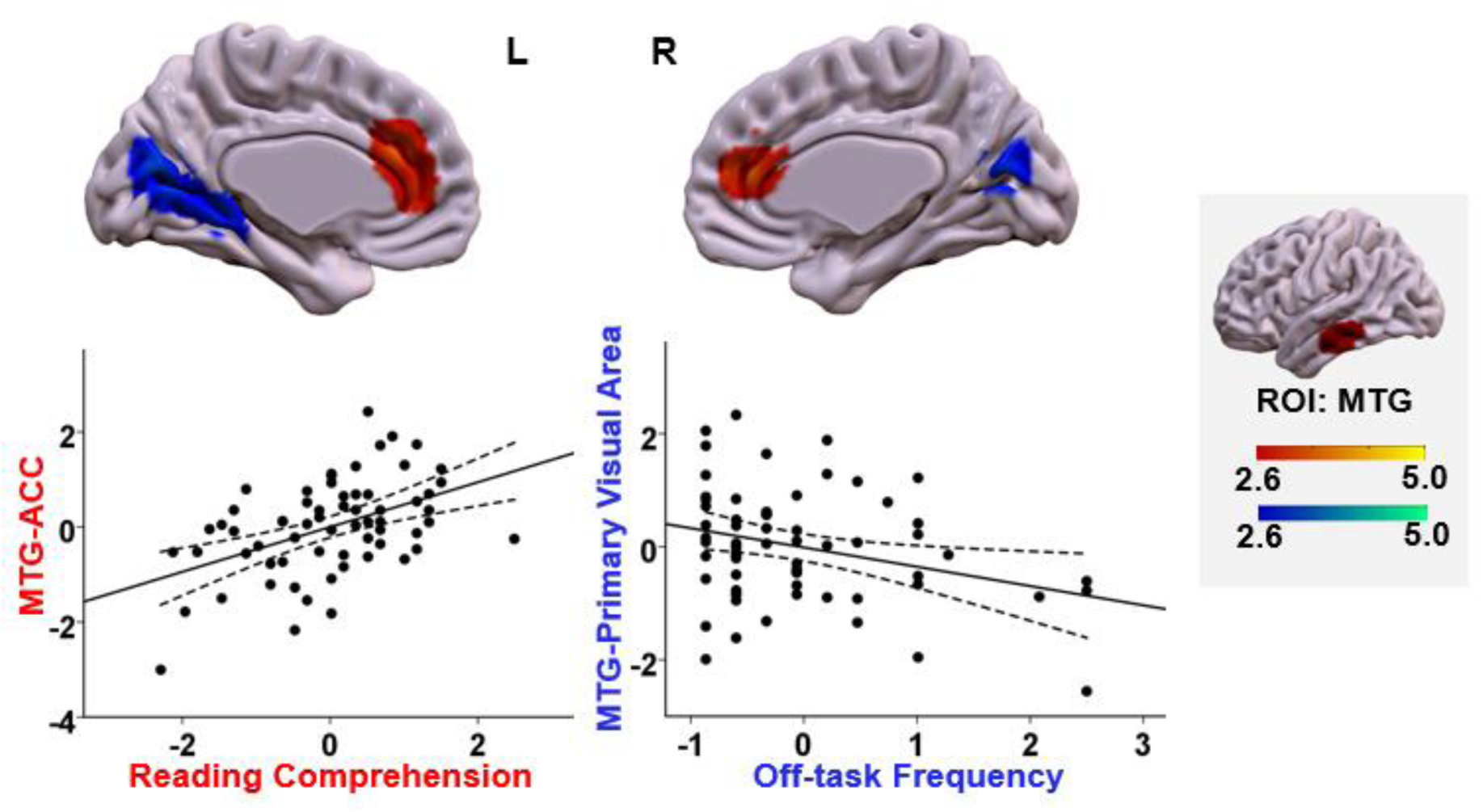
Functional connectivity of MTG linked to off-task frequency and comprehension. The regions in red showed stronger connectivity to MTG for participants with better comprehension, while the regions in blue showed weaker connectivity with MTG for participants with more frequent off-task thought. All maps are cluster corrected at a threshold of *z* > 2.6 (*p* < .05). The scatterplots present the correlation between behaviour (comprehension or off-task frequency) and the average correlation with the MTG seed and the relevant cluster (beta values). The error lines on the scatterplot indicate the 95% confidence estimates of the mean. Each point describes an individual participant. The right-hand panel highlighted in grey shows the seed region and the colour bars. MTG = middle temporal gyrus; ACC = anterior cingulate cortex.

#### 3.3.2. Off-task Frequency

We also found that increasing off-task frequency was associated with weaker connectivity between MTG in DMN and visual cortex (intracalcarine cortex, precuneus cortex, and lingual gyrus). This effect is presented in Figure 5. Of the voxels in this cluster that fell within one of the large-scale networks defined by Yeo et al. (2011), 100% were within the visual network. Consequently, participants with stronger intrinsic connectivity at rest between DMN and visual cortex were less likely to have lapses in attention to an ongoing reading task.

#### 3.3.3 Additional effects

Three further results failed to survive Bonferroni correction for the number of models (e.g., *p* < .0125). We do not seek to interpret these results but to guard against Type II errors, they are reported here.

i. With increasing off-task thought, a mid-cingulate somatomotor region showed greater disconnection with MTG (uncorrected *p* = .022; see Figure 6). This pattern resembles decoupling from visual cortex, found at a higher statistical threshold.
ii. Increasing off-task thought was associated with greater connectivity between left inferior frontal gyrus and MTG (uncorrected *p* = .044). Vatansever et al. (2017) similarly found poorer semantic performance with less segregation between temporal lobe DMN and frontal FPN regions at rest.
iii. Higher NYC-Q scores were linked to greater connectivity between MTG and parahippocampal gyrus as well as temporal fusiform cortex (uncorrected *p* = .028). Similarly, a recent fMRI study found stronger lateral temporal-to-hippocampal connectivity for people who tended to think about the past more during spontaneous thought (Smallwood et al., 2016). Although perceptual decoupling was related to the occurrence of off-task periods, hippocampal connectivity might be important for periods of off-task thought supported by episodic memory (Vincent et al., 2006).

**Fig. 6.**
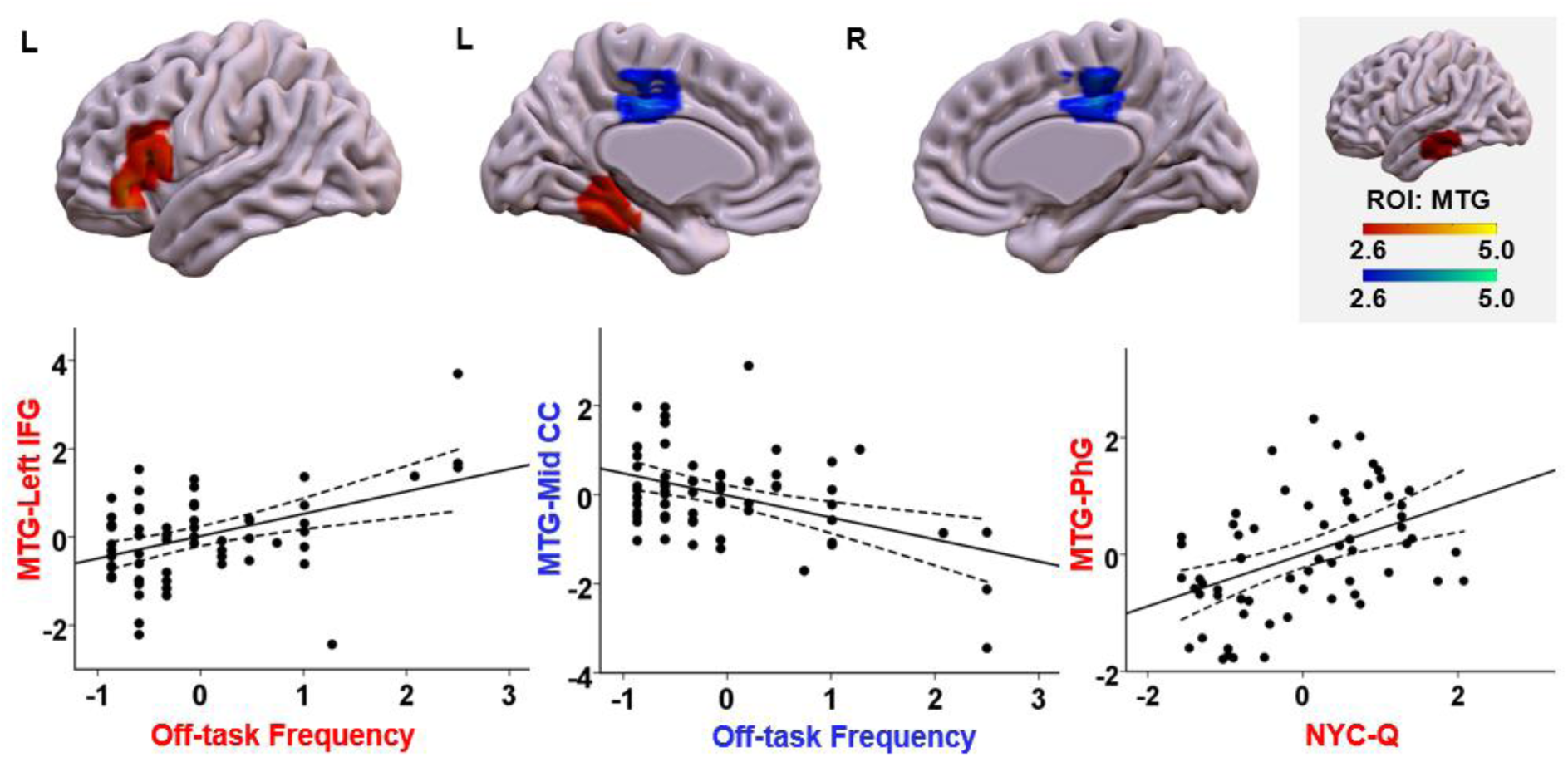
Additional effects of functional connectivity seeding from MTG. The region in warm colour indicates region displaying connectivity with MTG was stronger for the participants with frequent off-task thought or higher NYC-Q scores. And the regions in cold colour indicates regions displaying connectivity with MTG was weaker for the participants with frequent off-task thought. All maps are cluster corrected at a threshold of *z* > 2.6 (*p* < .05). The scatterplots present the relationship between the average correlation with MTG (beta values) in each region and off-task frequency, as well as NYC-Q scores. The error lines on the scatterplots indicate the 95% confidence estimates of the mean. Each point describes each participant. The upper right panel highlighted in grey shows the seed region and the colour bar. IFG = inferior frontal gyrus; Mid CC = middle cingulate cortex; PhG = parahippocampal gyrus; MTG = middle temporal gyrus.

In summary, consistent with the view that off-task thought is a perceptually-decoupled state, our analysis found that frequent off-task thought was associated with weaker connectivity between MTG within DMN and visual cortex. In addition, stronger connectivity between MTG and another DMN site in anterior cingulate cortex was associated with good understanding of the text. These findings show that connectivity within the DMN does not always predict off-task thought; instead, when MTG is strongly driven by visual input, participants tend to be better at comprehension of visually-presented texts, and when this region decouples from visual areas of the brain, people have greater frequency of off-task thoughts, which disrupts comprehension.

## 4. General discussion

We show how a region of the default mode network (DMN) within lateral temporal cortex (MTG) supports both on-task processing (i.e. comprehension) and off-task mental states (such as periods of off-task thought). Using an individual differences approach, we obtained support for the hypothesis that task-relevant patterns of cognition involve strong coupling to visual regions, while off-task cognition is more frequent in the face of perceptual decoupling. In Experiment 1, which employed a reading task in the scanner, we found people with good comprehension showed a greater BOLD response to orthographic inputs in both middle temporal gyrus (MTG) within DMN, and inferior temporal gyrus (ITG) in FPN. Since this effect was found for nonwords, it reflected a sensitivity to visual input in general, as opposed to activation that supported reading for meaning per se. In Experiment 2, we explored individual differences in the intrinsic connectivity of these sites. For individuals with more frequent off-task thought, MTG within DMN showed weaker connectivity with visual cortex, suggesting that perceptual decoupling may promote off-task thought. In contrast, for individuals with excellent comprehension, MTG showed greater connectivity with anterior cingulate cortex, also in the DMN. These findings show that DMN regions in lateral temporal cortex not only help us to understand incoming information, but they also build trains of thought that are independent of events in the environment. The behavioural trade-off between comprehension and off-task thought during reading may be explained by the sensitivity of temporal lobe DMN regions to visual inputs: when MTG is responsive to visual input, comprehension for visually-presented material is good, and when MTG is perceptually decoupled from visual cortex at rest, participants report being off-task more frequently.

In this way, our results suggest that MTG contributes to different mental states through distinct patterns of functional connectivity: rather than different temporal lobe regions supporting on-task and off-task semantic retrieval (for example, regions falling within FPN and DMN respectively), diverse patterns of connectivity from the same DMN region can underpin both off-task thought and comprehension. It has already been observed that semantic regions within temporal cortex have a pattern of connectivity to both DMN core and visual cortex (Binney, Embleton, Jefferies, Parker, & Lambon Ralph, 2010; Murphy et al., 2017; Visser, Jefferies, Embleton, & Lambon Ralph, 2012) – our results suggest that both of these connections are important for good comprehension during a reading task. Our results are also broadly consistent with the recent observation that while DMN often shows a response to nonwords over words (reflecting off-task processing), these DMN regions also classify the meaningfulness of input (Mattheiss, Levinson, & Graves, 2018).

Functionally, MTG is implicated in heteromodal aspects of cognition as the inputs along auditory and visual processing streams maximally converge here (see also Lambon Ralph et al., 2017; Margulies et al., 2016; Murphy et al., 2017; Visser et al., 2012). MTG responds more strongly to memory-based and meaning-based decisions, consistent with the location of this cluster at the anterior end of the ventral visual stream and yet also within the DMN (Murphy et al., 2018). In line with these studies, the anterior and middle temporal lobe have been identified as important for text comprehension (Ferstl, Neumann, Bogler, & Von Cramon, 2008; Jangraw et al., 2018; Kuperberg, Lakshmanan, Caplan, & Holcomb, 2006). Nevertheless, activation within MTG is insufficient for comprehension – our results suggest this region also needs to be strongly activated by visual inputs and to interact with other regions of DMN implicated in comprehension.

Consistent with the view that off-task thought is a perceptually-decoupled state (Smallwood, 2013), our analysis revealed that frequent off-task thought was associated with weaker correlation between MTG and primary visual cortex. Event related potentials evoked by sensory inputs are reduced in magnitude during episodes of off-task thought, relative to on-task periods (Kam et al., 2011). The posterior core of the DMN which supports heteromodal integration (Braga, Sharp, Leeson, Wise, & Leech, 2013) also contributes to different types of spontaneous thought (Smallwood et al., 2016). The role of MTG in comprehension and off-task thought may be similar: perceptual decoupling of MTG from visual cortex may allow this region to support off-task thought that is unrelated to the immediate external environment. By this view, the occurrence of off-task thought does not always reflect the failure of high-level attention or self-monitoring systems; instead the absence of perceptual input disrupts performance on externally-presented tasks, and in this way, relatively low-level processes contribute to higher-order cognitive states.

While DMN regions may support off-task states which impair comprehension, connectivity within DMN predicted good comprehension in the current study. In people with better comprehension, MTG coupled more with anterior cingulate cortex: both the MTG seed and much of the ACC cluster fell largely within DMN as defined by Yeo et al. (2011). Both of these regions have been previously argued to form ‘hubs’ that integrate diverse elements of cognition (Margulies et al., 2016; Rossell, Bullmore, Williams, & David, 2001; Visser et al., 2012; Zhao et al., 2017). ACC shows graded connections at rest with both sensory and motor cortices, as well as with memory/DMN regions (Margulies et al., 2007). Consequently, participants with strong connectivity between these ‘hubs’ might have more efficient retrieval of conceptual information, and this would improve reading comprehension when visual inputs penetrate DMN. Although there are likely to be functional subdivisions within DMN, these findings rule out the hypothesis that DMN regions are always off-task and that their activation/connectivity is necessarily linked to poor performance on ongoing tasks, contradicting a commonly-proposed view (Mazoyer et al., 2001; Raichle et al., 2001; Shulman et al., 1997).

In conclusion, we found that dissociable patterns of activation and intrinsic connectivity in MTG within DMN predicted comprehension and off-task thought. Better comprehension was associated with greater coupling of MTG with another DMN region in anterior cingulate gyrus. In contrast, greater disconnection between MTG and primary visual cortex was associated with frequent off-task thought. We conclude that DMN regions in lateral temporal cortex not only help us to understand information in the external environment, but also form thoughts that are independent from what is happening around us – however, both of these aspects of cognition are also supported by a broader network of brain regions. By examining individual differences in comprehension and off-task thought as opposed to task activation (Seghier & Price, 2018), we were able to establish which connectivity differences are functionally predictive. Differences in the responsiveness and intrinsic architecture of the pathway from vision, via MTG to DMN predicted both comprehension and the frequency of off-task thought during reading.

## Acknowledgements

We would like to thank Tirso Gonzalez Alam, Theodoros Karapanagiotidis and Charlotte Murphy for their help with data analysis. EJ was supported by the European Research Council (Project ID: 771863 - FLEXSEM), JS was supported by the European Research Council (Project ID: 646927- WANDERINGMINDS), and MZ was supported by the China Scholarship Council (CSC) Scholarship (No. 201704910952).

